# Human rod photoreceptor outer segments are supported by accessory inner segment structures

**DOI:** 10.1101/2024.08.09.607370

**Authors:** Tylor R. Lewis, Natalia V. Klementieva, Sebastien Phan, Carson M. Castillo, Keun-Young Kim, Lauren Y. Cao, Mark H. Ellisman, Vadim Y. Arshavsky, Oleg Alekseev

## Abstract

The first steps in vision take place in photoreceptor cells, which are highly compartmentalized neurons exhibiting significant structural variation across species. The light-sensitive ciliary compartment, called the outer segment, is located atop of the cell soma, called the inner segment. In this study, we present an ultrastructural analysis of human photoreceptors, which reveals that, in contrast to this classic arrangement, the inner segment of human rods extends alongside the outer segment to form a structure hereby termed the “accessory inner segment”. While reminiscent of the actin-based microvilli known as “calyceal processes” observed in other species, the accessory inner segment is a unique structure: (1) it contains an extensive microtubule-based cytoskeleton, (2) it extends far alongside the outer segment, (3) its diameter is comparable to that of the outer segment, (4) it contains numerous mitochondria, and (5) it forms electron-dense structures that likely mediate adhesion to the outer segment. Given that the spacing of extrafoveal human photoreceptors is more sparse than in non-primate species, with vast amounts of interphotoreceptor matrix present between cells, the closely apposed accessory inner segment likely provides structural support to the outer segment. This discovery expands our understanding of the human retina and directs future studies of human photoreceptor function in health and disease.

## Introduction

Light detection confers a substantial evolutionary advantage to all life forms, from cyanobacteria to humans. In most species, it is accomplished by highly compartmentalized photoreceptor neurons, which initially developed in early organisms more than 600 million years ago (Lamb et al., 2007). Photoreceptors contain specialized cellular compartments highly enriched in visual pigments and downstream signaling proteins, thereby achieving remarkable light sensitivity, including single photon detection (Baylor et al., 1979).

In vertebrate photoreceptors, the light-sensitive compartment, called the outer segment (OS), is a modified primary cilium consisting of hundreds of flattened membrane discs enriched in visual pigments. The OS emanates from the apical end of the photoreceptor cell body, called the inner segment (IS), which performs housekeeping functions of the cell. At the opposite end of the photoreceptor cell is located the synaptic terminal, which propagates visual signals to downstream neurons. This general anatomical organization has been described for a wide variety of species (reviewed in (Goldberg et al., 2016; Spencer et al., 2020; Wensel et al., 2016)), including humans (e.g., (Ahnelt et al., 1987; Bairati and Orzalesi, 1963; Cohen, 1965; Missotten, 1960; Villegas, 1964)).

In this study, we revisited the ultrastructure of human photoreceptors and uncovered a unique anatomical feature overlooked in previous studies. We demonstrate that, in contrast to other well-characterized vertebrate species, the IS of human rod photoreceptors forms a microtubule-based extension positioned alongside the OS. This structure, hereby termed the “accessory inner segment” (aIS), contains electron-dense elements spanning the aIS and OS plasma membranes, which likely mediate their adhesion. Given that the spacing of individual human rods is significantly sparser than what is reported for non-primate species, a likely function of the aIS may be to enhance the structural stability of the OS.

## Results and Discussion

### Human rod photoreceptors have an accessory inner segment

Ultrastructural analysis of longitudinally sectioned healthy human retinas using transmission electron microscopy (TEM) revealed an immediate striking observation that sets human rod photoreceptors apart from those of essentially all other previously characterized vertebrate species. In other species, there is a clear vertical distinction between the photoreceptor IS and OS, with the OS located directly apical to the IS (Pearring et al., 2013) (**Figure 1A**). In contrast, a large portion of the IS in human rods extends alongside the OS to form structures we have termed “accessory inner segment” (aIS; **Figure 1B**). The intracellular contents of the aIS appear to be comparable to the rest of the IS, including the presence of multiple mitochondria and long filamentous structures.

**Figure 1.**
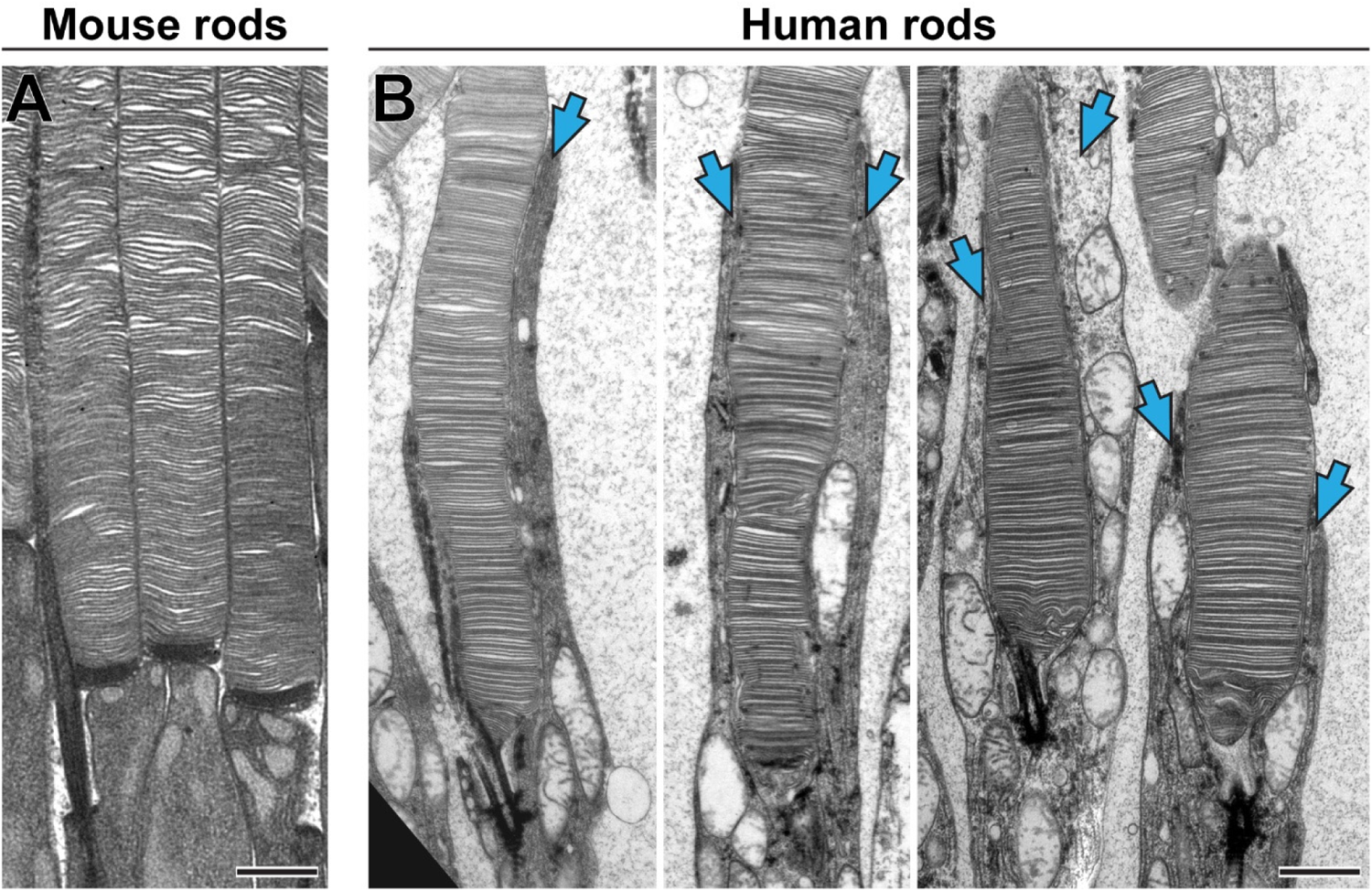
Human rod photoreceptors have an accessory inner segment. TEM images of rod photoreceptors from longitudinally sectioned mouse (A) and human (B) retinas. Blue arrows indicate aIS. Scale bars: 1 µm

To further investigate the structure and composition of the aIS, we performed ultrastructural analysis of tangentially sectioned human retinas. We observed that rod OS are typically accompanied by adjacent aIS (**Figure 2A**). At the base of the OS, the aIS can be as wide, if not wider, as the OS and contains several mitochondria (Example 1 in **Figure 2A**). As it extends along the OS, the aIS narrows (Example 2) and eventually becomes devoid of mitochondria (Example 3). At higher magnification, the filamentous material within aIS was identified to be densely arranged microtubules (**Figure 2B**). This was further corroborated by immunostaining of human rods with antibodies against β-tubulin (**Figure 2C**), which revealed two microtubular structures emanating from the distal end of each rod IS, one corresponding to the ciliary axoneme and the other to the aIS. This is in contrast to the pattern of tubulin staining in mammalian species lacking the aIS, which yields staining of only the ciliary axoneme (e.g., (Bosch Grau et al., 2017; Sahly et al., 2012)).

**Figure 2.**
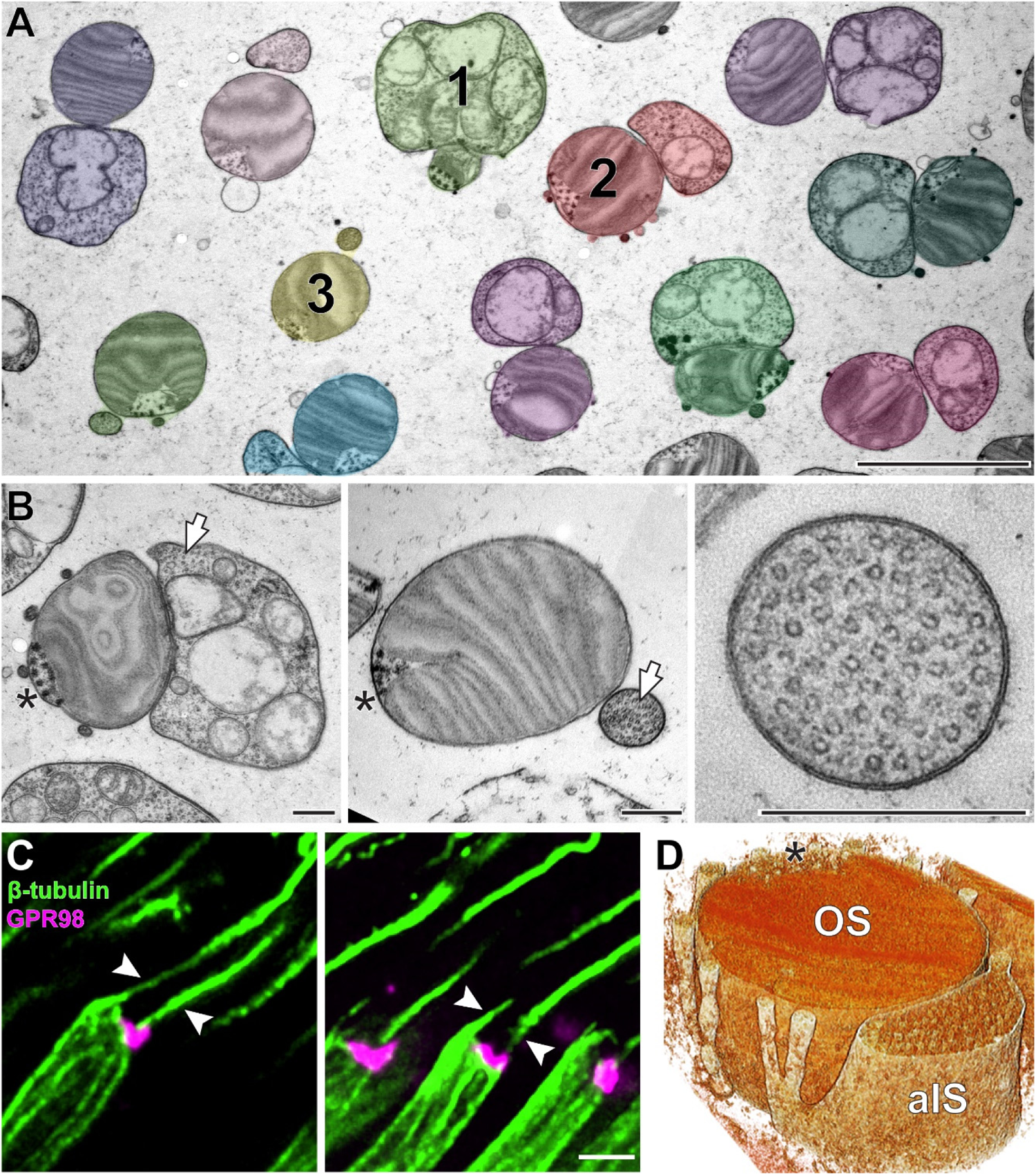
The accessory inner segment contains microtubules. **(A)** A low-magnification TEM image of tangentially sectioned human retina. Rods containing a paired aIS and OS are pseudo-colored. Three numbered cells are sectioned at different axial positions ranging from proximal (example 1) to intermediate (example 2) and distal (example 3). **(B)** TEM images of human rods tangentially sectioned at intermediate (left panel) and distal (middle panel) positions. White arrows indicate bundles of microtubules inside the aIS. The right panel shows a magnified image of the distal aIS devoid of mitochondria and tightly packed with microtubules. **(C)** Immunofluorescence images of rods from longitudinally sectioned human retina, in which microtubules (green) and the connecting cilium region (magenta) are labeled with antibodies against β-tubulin and GPR98, respectively. Arrowheads indicate microtubules emanating from opposite sides of the OS base within an individual rod. **(D)** Rendering of the 3D-ET shown in **Supplementary Video 1**. Asterisks in (B) and (D) indicate the ciliary axoneme. Scale bars: 5 µm (A); 0.4 µm (B); 2 µm (C).

To better visualize the architecture of the aIS, we employed 3-dimensional electron tomography (3D-ET). Representative tomograms of two rods are shown in **Supplementary Videos 1 and 2,** and a rendering of the tomogram from **Supplementary Video 1** spanning ∼600 nm of the axial photoreceptor length is shown in **Figure 2D**. 3D-ET revealed another notable feature of aIS organization – the presence of smaller microtubule-containing processes branching off of the main aIS structure.

In some ways, the aIS and its branches resemble calyceal processes, which are microvilli-like extensions emanating from the IS and surrounding the OS in many species (reviewed in (Goldberg et al., 2016)). However, calyceal processes are supported by an actin network, not microtubules (e.g., (Sahly et al., 2012)), indicating that aIS are not simply large calyceal processes but rather a previously unrecognized type of structure observed thus far only in human rods. In fact, smaller processes that appear more similar to traditional calyceal processes were observed in human rods (**Figure 3A-C**). They emanate from the distal end of the rod IS in the area behind the connecting cilium, which we reconstructed using 3D-ET (**Figure 3D** and **Supplementary Video 3**).

**Figure 3.**
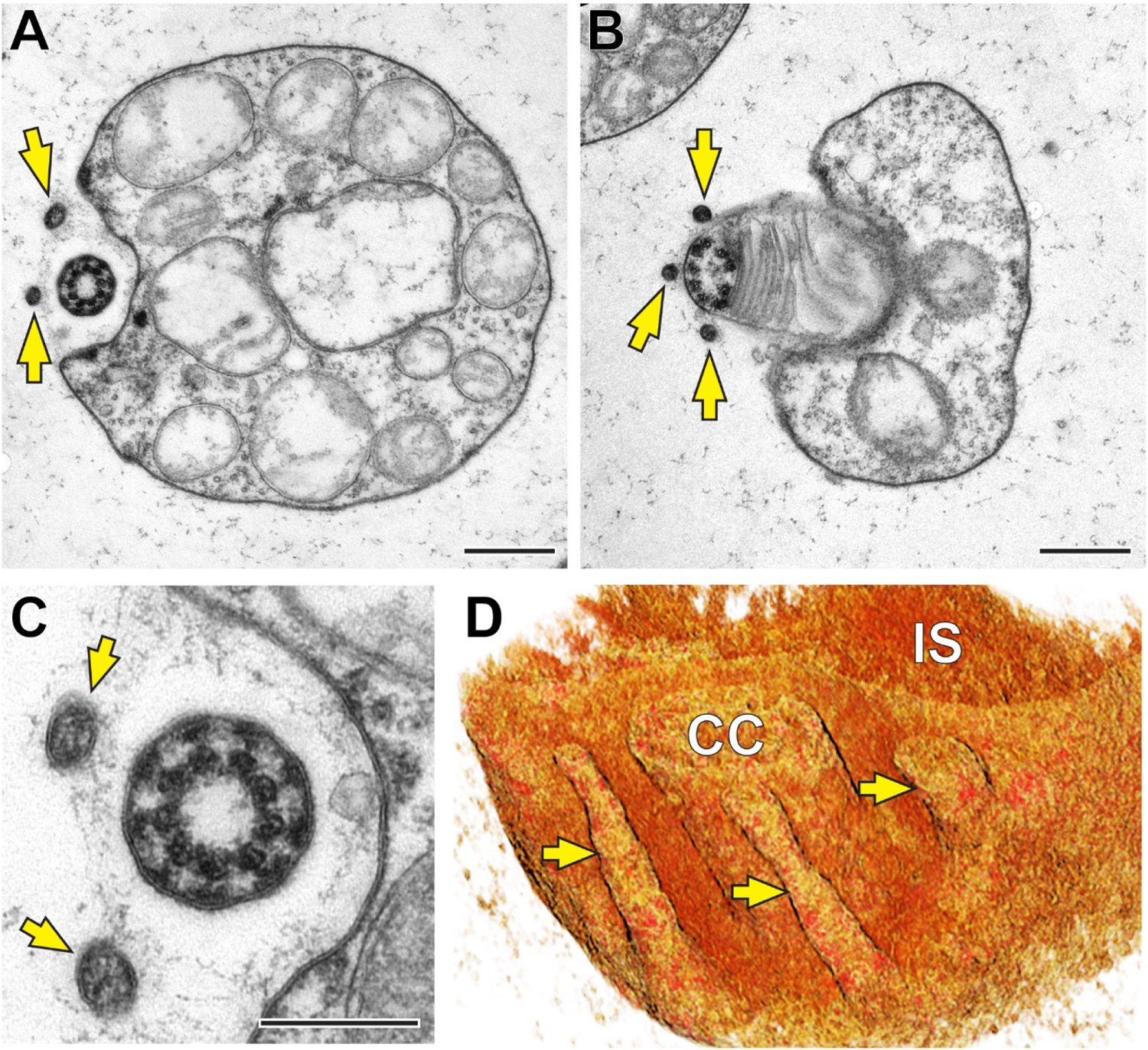
Presumed calyceal processes are located in the ciliary region of human rods. **(A-C)** TEM images of rods from tangentially sectioned human retina taken at the level of the connecting cilium (CC). **(D)** Rendering of the 3D-ET shown in **Supplementary Video 3**. In all panels, yellow arrows indicate presumed calyceal processes. Scale bars: 0.5 µm (A,B); 0.25 µm (C).

In contrast to rods, we did not encounter the presence of aIS in human cones (**Figure 4**). Instead, proximal regions of cone OS are surrounded only by multiple thin microvilli-like extensions previously described as calyceal processes (e.g., (Sahly et al., 2012)).

**Figure 4.**
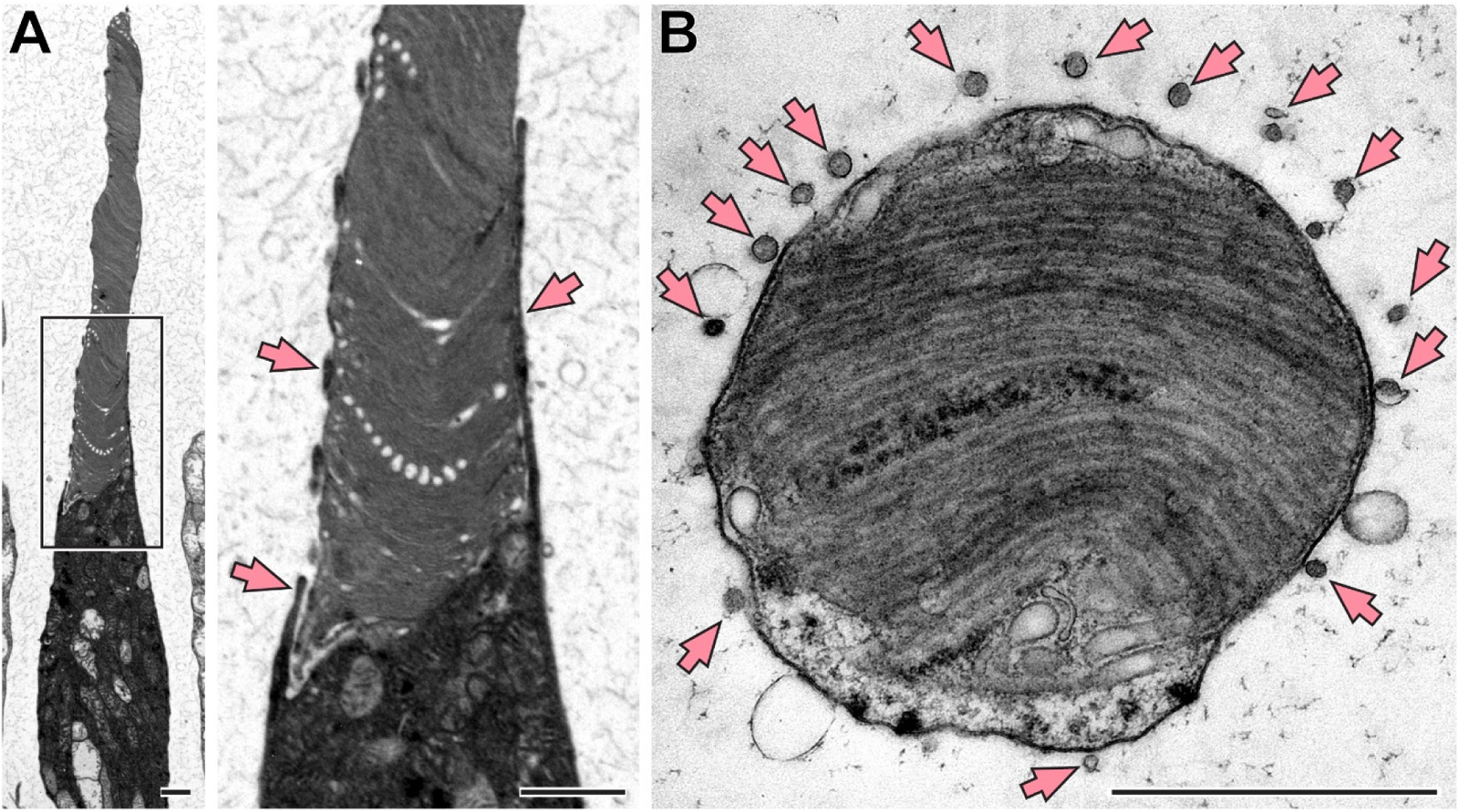
Human cone photoreceptors do not have an accessory inner segment TEM images of cones from longitudinally. **(A)** and tangentially **(B)** sectioned human retinas. The boxed area in the left panel is magnified in the middle panel. Pink arrows indicate calyceal processes surrounding a cone OS. Scale bars: 1 µm.

Overall, we conclude that the aIS is a unique feature of human rods distinct from calyceal processes. In the next part of this study, we turned our attention to investigating the possible role of this unusual structure.

### The accessory inner segment is connected to the outer segment via electron-dense structures

The functional role of the aIS is not immediately clear. One consideration is that the spacing of human extrafoveal photoreceptors in the IS/OS region is sparse relative to many non-primate species, including mice (e.g., (Lindell et al., 2023)). This sparse organization can be visualized by imaging human retinas tangentially sectioned at different planes, ranging from the outer limiting membrane (OLM; located at the proximal base of the IS) to the IS/OS junction (**Figure 5A**). Around the OLM, individual human photoreceptor cells are fully separated by Müller glial cells with which they form adherens junctions. The only connections between adjacent photoreceptors consist of gap junctions documented in previous studies ((Cohen, 1989; Uga et al., 1970); **Supplementary Figure 1**). Immediately above the OLM, endfeet microvilli of Müller glial cells occupy most of the space between photoreceptors. At the OS base, photoreceptor cells remain separated to a similar degree by an abundant amount of interphotoreceptor matrix material. In contrast, mouse photoreceptors are densely packed at all levels (**Figure 5A**).

**Figure 5.**
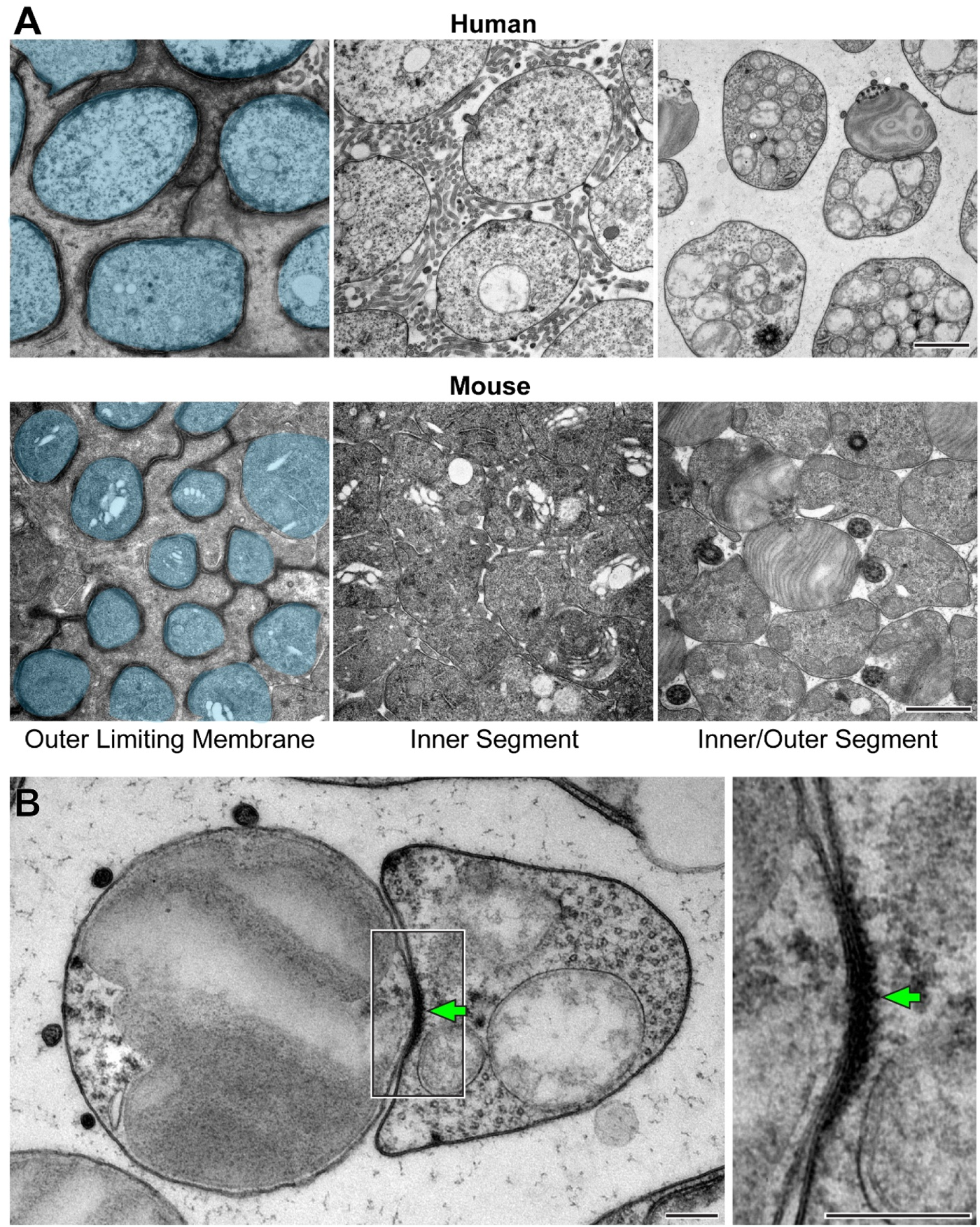
The accessory inner segment is connected to the outer segment via electron-dense structures. **(A)** TEM images of human and mouse retinas tangentially sectioned at different levels, as indicated. Photoreceptor cells in the left panels are pseudo-colored blue to distinguish them from Müller glial cells. **(B)** TEM image of a rod from tangentially sectioned human retina. The boxed area on the left is magnified on the right. Green arrows indicate an electron-dense structure located between aIS and OS. Scale bars: 1 µm (A); 0.2 µm (B).

Considering this spatial arrangement of human rods, we speculate that the functional role of the aIS is to provide physical support to the sparsely spaced OS. This hypothesis presumes that the OS and the adjacent aIS are not merely juxtaposed but also physically connected.Therefore, we sought to investigate whether there are any structural features at their interface. Using TEM and 3D-ET, we identified electron-dense structures that span the plasma membranes of OS and aIS and bring these membranes remarkably close to one another (**Figure 5B** and **Supplementary Video 4**). These electron-dense structures could serve to maintain the close apposition between aIS and OS, which would be necessary for the aIS to provide physical support to the OS.

### Human rod discs have multiple incisures despite having a small diameter

The disc membranes stacked inside the rod OS have indentations in their rims, called incisures. The number of incisures in each disc varies across species (Cohen, 1960; Nilsson, 1965; Sjostrand, 1953). For example, the relatively small-diameter discs of mouse rods have only a single incisure, whereas the large diameter discs of frog rods have up to several dozen (e.g., (Lewis et al., 2023)). It has been generally assumed that the larger the disc diameter is, the greater is the number of incisures (Tsukamoto, 1987). Contrary to this notion, human rods, which have a relatively small disc diameter, have been reported to have numerous incisures (Cohen, 1965).

We revisited the relationship between the disc diameter and the number of incisures by analyzing rods from four species – humans, mice, *Xenopus* frogs and zebrafish – and found no correlation (**Figure 6**). Despite having a comparable diameter to mouse rod discs, human rod discs have several incisures, much like the large-diameter discs of frog rods. Furthermore, the large-diameter zebrafish discs have only a single incisure. These observations refute the notion that the number and complexity of incisures correlates with disc diameter. Whether the variability in incisure number bears any biological significance remains unknown.

**Figure 6.**
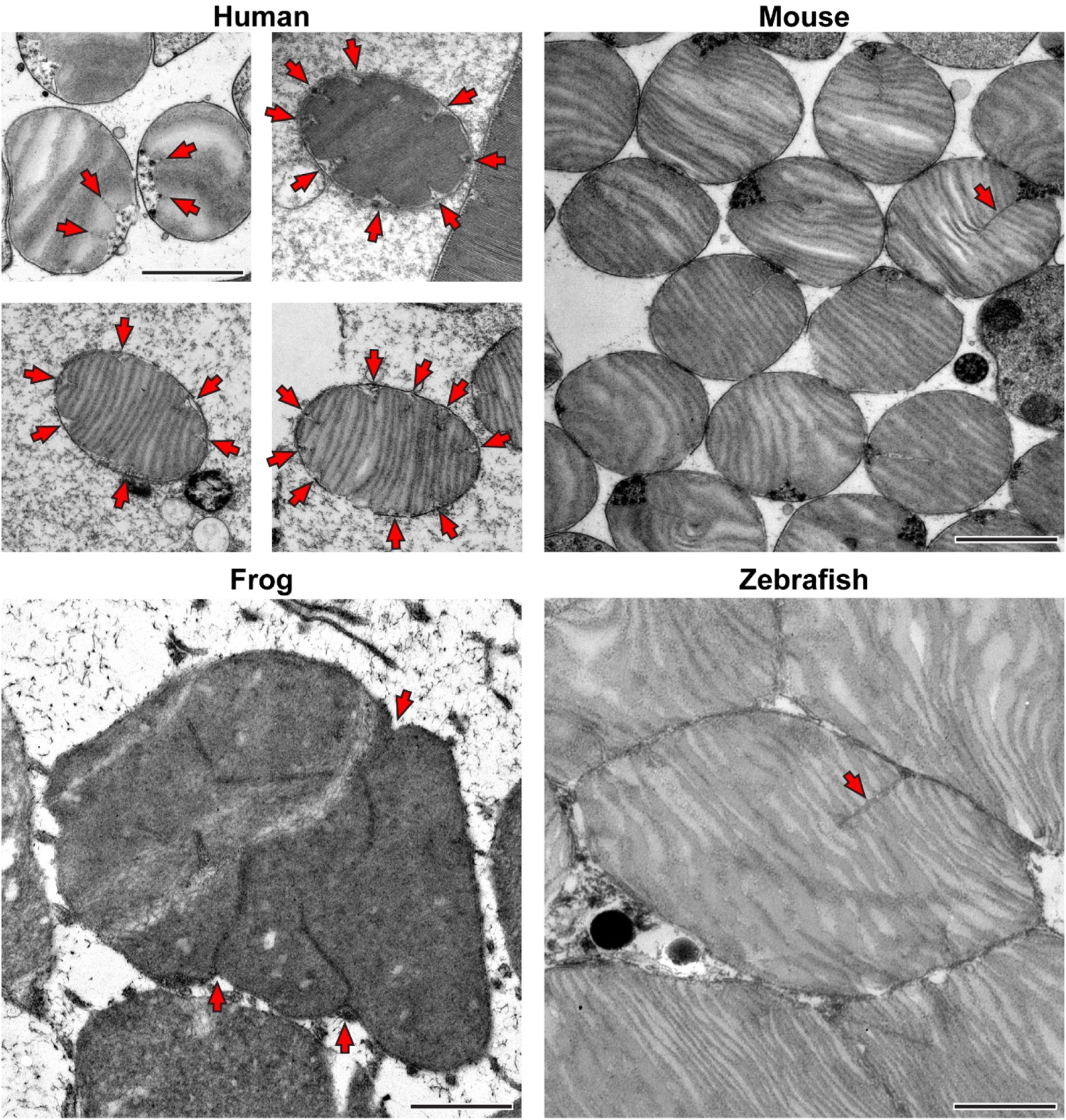
Human rod discs have multiple incisures. TEM images of rods from human, mouse, frog and zebrafish tangentially sectioned retinas. Red arrows indicate disc incisures. Scale bars: 1 µm.

Our analysis of human rods revealed several further observations about the organization of incisures. At the OS base, we typically observed two shallow incisures, each originating next to the ciliary axoneme (**Figure 6**). More distally in the OS, the number of incisures increases to as many as 10 per disc. Of note, 3D-ET revealed that incisures located in sequential discs within a stack are not well-aligned across discs (**Supplementary Video 1** and **Supplementary Figure 2**). Similar to our previous findings in mice (Lewis et al., 2023), many incisures contained electron-dense structures at their terminal ends, as well as structures that appear to connect the two apposing sides of an incisure (**Supplementary Figure 3**). These structural elements may initiate the formation of an incisure and/or maintain its integrity.

### Human photoreceptors release extracellular vesicles from their inner segments

We have recently shown that in several non-primate species, photoreceptors release extracellular vesicles from their IS (Lewis et al., 2022), which can serve as a means of disposing of undesirable material (Lewis et al., 2024). We now show that the same phenomenon takes place in the human retina. Several examples of extracellular vesicles that were captured either during or immediately following their budding off of the IS are shown in **Figure 7**. This finding highlights the general nature of this phenomenon and opens exciting questions about the role of vesicular release in maintaining the health of photoreceptor cells.

**Figure 7.**
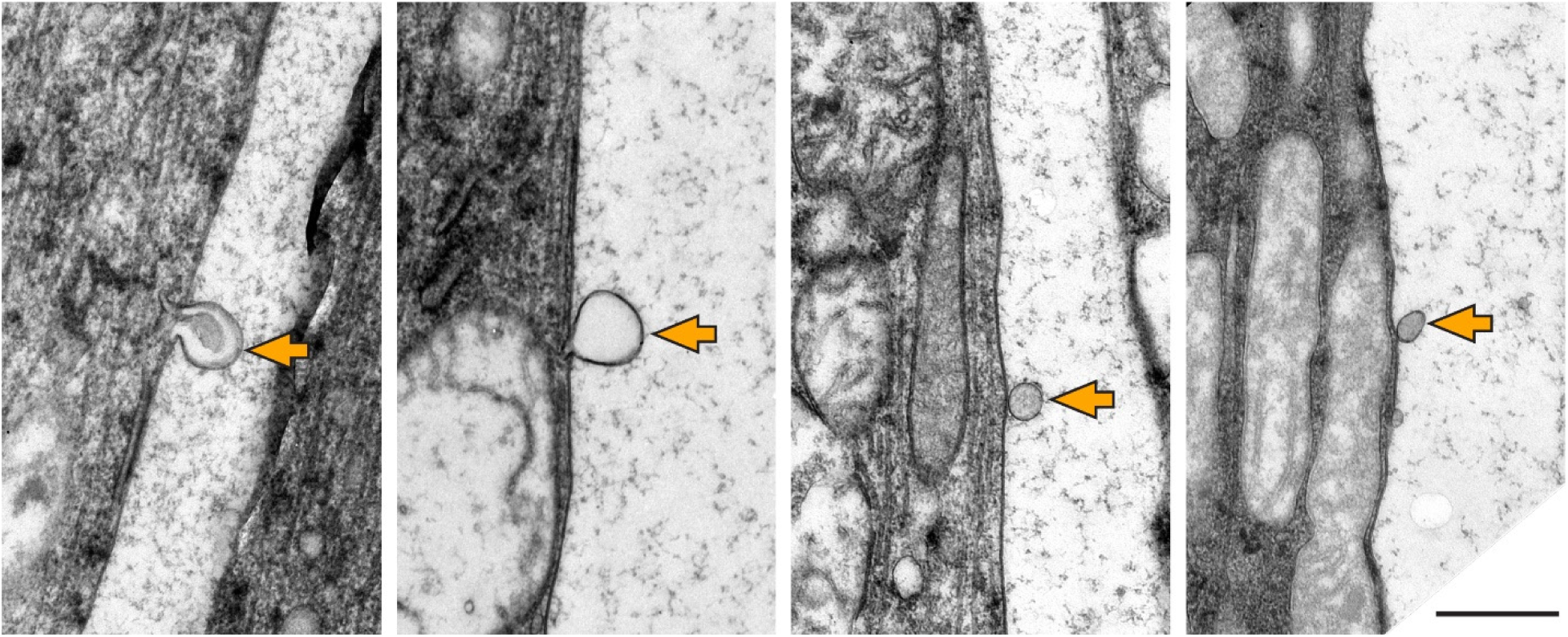
Human photoreceptors release extracellular vesicles from their inner segment. TEM images of photoreceptors from longitudinally sectioned human retina. Orange arrows indicate extracellular vesicles captured either in the process of or immediately following release from the IS. Scale bar: 0.5 µm.

### Concluding remarks

The central finding of this study is that human rod photoreceptors possess a unique structure consisting of a large protrusion of the IS reinforced by an extensive microtubule-based network and connected to the OS through distinct intermembrane adhesions. A schematic illustration of this “accessory inner segment” (aIS) is shown in **Figure 8**. To the best of our knowledge, the presence of this structure has not been reported in either humans or other species, including the thoroughly characterized macaque monkeys (Sahly et al., 2012). Whether aIS are present in rods of other non-human primates, particularly apes, is an interesting evolutionary question warranting future studies.

**Figure 8.**
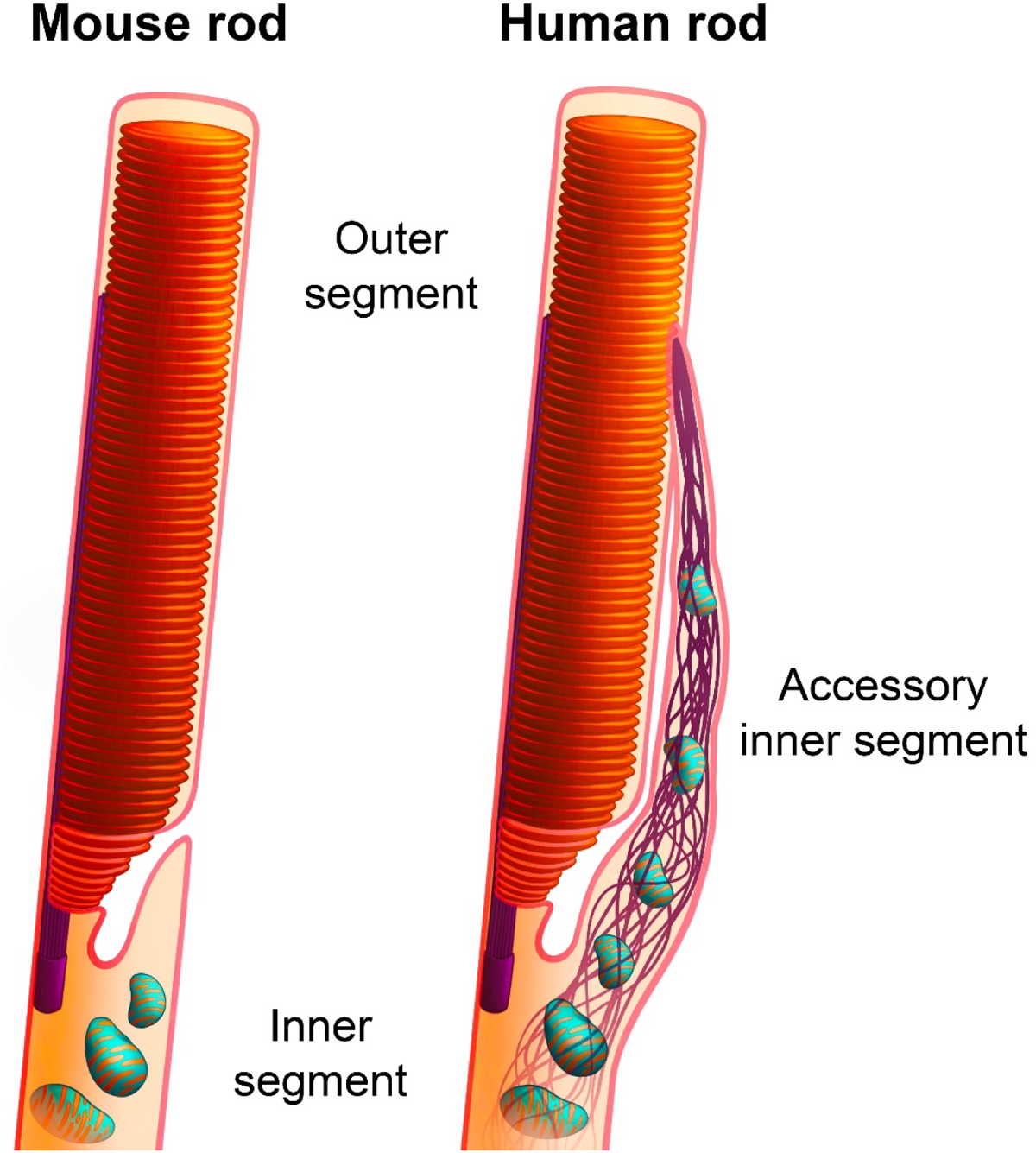
Schematic representation of the accessory inner segment of human rod photoreceptors. The image on the left represents the typical vertical arrangement of the IS and OS in most studied species, such as the mouse. The image on the right illustrates the new observation from this study that human rods have an aIS extending from the IS alongside the OS. The base of the aIS is comparable in width to the OS and contains mitochondria as well as a microtubule-based cytoskeleton. The apical portion of the aIS tapers in diameter and contains a dense bundle of microtubules. For ease of visualization, the smaller microtubule-containing processes branching off of the main aIS structure, as well as inner segment microtubules, are not shown.

While it is conceivable that aIS merely fill the space between the sparsely packed human photoreceptors, the presence of unique electron-dense structures juxtaposing the membranes of the OS and the aIS argues otherwise. The fact that the aIS contains an extensive microtubule network further suggests that the most plausible function of the aIS is to provide mechanical support to the OS. The significance of such support remains to be determined, as well as whether a disruption of this structure or loss of its association with the OS may affect normal photoreceptor function. It is well-documented that many hereditary visual disorders affecting human photoreceptors are either not phenocopied in mice bearing the same mutations (e.g., (Yang et al., 2012)) or the respective genes are not even present in mice (e.g., (Abd El-Aziz et al., 2008)). It is tempting to speculate that at least some of these differences are explained by the lack of the aIS in photoreceptors of commonly used animal models.

## Methods

### Human donor eye tissue

Enucleated human globes were obtained by Miracles in Sight (Winston-Salem, NC) and distributed by the BioSight Tissue Repository and Service Center (Duke University) under an approved Institutional Review Board protocol. Retinas used for electron microscopy analysis originated from a 61-year-old male and a 73-year-old female, whereas retinas used for immunofluorescence microscopy originated from a 63-year-old female. Only donors without medical history of ocular pathology were selected. All tissues were de-identified prior to being delivered to the laboratory. Whole globes were enucleated and kept on ice in a humidified chamber for 6-12 hours until further processing.

### Animal husbandry

Animal maintenance and experiments were approved by the Institutional Animal Care and Use Committees at Duke. WT mice (*Mus musculus*) were C57BL/6J (Jackson Labs stock #000664). WT zebrafish (*Danio rerio*) were AB line (ZFIN stock #ZDB-GENO-960809-7). Images of WT frogs (*Xenopus tropicalis*) were obtained during the course of a previous collaborative study with the National Xenopus Resource (Lewis et al., 2023) and used with permission of Marko Horb (Marine Biological Laboratory). All experiments were performed with animals of randomized sex.

### Transmission electron microscopy (TEM)

For humans, enucleated globes were fixed 6-7 hours after death in a solution of 2% paraformaldehyde, 2% glutaraldehyde and 0.05% calcium chloride in 50 mM MOPS (pH 7.4) for 1 week. Following the removal of the anterior segment and vitreous humor, 12 mm punches of retina were obtained. For longitudinal sections, tissue was embedded in 2.5% low melting point agarose (Precisionary) and cut into 200 µm thick slices on a Vibratome (VT1200S; Leica). For tangential sections, dissected tissue was processed whole.

For mice, anesthetized animals were transcardially perfused with a solution of 2% paraformaldehyde, 2% glutaraldehyde and 0.05% calcium chloride in 50 mM MOPS (pH 7.4). Enucleated eyes were fixed for an additional 2 hours in the same fixation solution at RT. Following the removal of the cornea and lens, agarose sections from eyecups were cut as described above.

For zebrafish, enucleated eyes were fixed with 2% paraformaldehyde and 2% glutaraldehyde in 0.1 M sodium cacodylate buffer (pH 7.2) overnight at 4°C. Following the removal of the cornea and lens, agarose sections from eyecups were cut as described above.

Resulting agarose sections or tissues were stained with 1% tannic acid (Electron Microscopy Sciences) and 1% uranyl acetate (Electron Microscopy Sciences), gradually dehydrated with ethanol and infiltrated and embedded in Spurr’s resin (Electron Microscopy Sciences). 70 nm sections were cut, placed on copper grids and counterstained with 2% uranyl acetate and 3.5% lead citrate (19314; Ted Pella). The samples were imaged on a JEM-1400 electron microscope (JEOL) at 60 kV with a digital camera (BioSprint; AMT). Image analysis and processing was performed with ImageJ.

The quality of tissue preservation of human eyes was assessed prior to conducting ultrastructural analysis by examining 500 nm thick sections stained with methylene blue (**Supplementary Figure 4A**). Images were acquired using a confocal microscope (Eclipse 90i and A1 confocal scanner; Nikon) with a 60× objective (1.49 NA Plan Apochromat VC; Nikon) and NIS-Elements software (Nikon).

### 3-Dimensional intermediate high voltage electron microscopic tomography (3D-ET) of thick sections

Either 250 or 750 nm thick retinal sections were cut and placed on 50 nm Luxel film slot grids. Grids were glow-discharged on both sides, and a mixture of 10 nm, 20 nm and 60 nm gold particles were deposited on the sample surfaces to serve as fiducial markers. 3D-ET was conducted on a Titan Halo (FEI, Hillsboro, OR, USA) operating at 300 kV in TEM mode for 250 nm sections or in STEM mode for 750 nm sections. A four-tilt series data acquisition scheme previously described (Phan et al., 2017) was followed in which the specimen was tilted from −60° to +60° every 0.25° at four evenly distributed azimuthal angle positions. Images were collected on an 8k×8k direct detector (DE64; Direct Electron, San Diego, CA, USA) in TEM mode or with a high-angle annular dark field (HAADF) detector in STEM mode. The final volumes were generated using an iterative reconstruction procedure (Phan et al., 2017). 3dmod and ImageJ were used for image analysis, and Amira was used for volume rendering.

### Immunofluorescence microscopy

Enucleated human globes were dissected 12 hours after death. Following the removal of the anterior segment and vitreous humor, the posterior segment was cut into several pieces, and retinal tissue carefully removed and fixed in 4% paraformaldehyde in PBS for 1 hr. Tissue was embedded in low melting point agarose and sectioned using a Vibratome at 100 µm thickness. Sections were blocked in PBS containing 7% donkey serum and 0.5% Triton X-100 for 30 min at RT and incubated with primary antibodies against β-tubulin (1:500; T4026; Millipore-Sigma) and GPR98 (1:200; PA5-84761; Thermo Fisher Scientific) overnight at 4°C. Sections were washed and incubated with secondary donkey anti-mouse and anti-rabbit antibodies conjugated to Alexa Fluor 488 or 568 (1:1000; A21202 and A10042, respectively; Thermo Fisher Scientific) for 2 hr at RT. Sections were washed and mounted onto slides with Shandon Immu-Mount (9990402; Thermo Fisher Scientific). Images were acquired using a confocal microscope (Eclipse 90i and A1 confocal scanner; Nikon) with a 60× objective (1.49 NA Plan Apochromat VC; Nikon) and NIS-Elements software (Nikon). Optical *z*-sections were collected at a resolution of 1024 × 1024 pixels with a step size of 0.15 μm. 3D deconvolution was performed in NIS-Elements, and deconvolved *z*-stacks were processed using ImageJ. Images are shown as maximum intensity *z*-projections over depth of 1.5 μm. The quality of tissue preservation was assessed by immunofluorescence staining with Höechst 33342 (10 µg/mL; 62249; Thermo Fisher Scientific) and fluorescent-conjugated Wheat Germ Agglutinin (WGA) (1 µg/mL; W11261; Thermo Fisher Scientific) (**Supplementary Figure 4B**).

## Acknowledgements

The authors would like to thank Marko Horb (National Xenopus Resource at the Marine Biological Laboratory) for allowing us to use an image obtained during the course of our previous study as a cross-species comparison, and Andrew Alekseev for generating the artwork. This work was supported by the National Institutes of Health grants EY033763 (TRL), EY030451 (VYA), EY005722 (VYA), EY033857 (OA), Duke University Physician-Scientist Strong Start Award (OA), Unrestricted Award from Research to Prevent Blindness Inc. (Duke University), as well as grants to MHE for NCMIR facilities, staff and brain research: NSF 2014862, R01AG081037, U24NS120055, S10OD021784 and R01GM138780.

---

**Supplementary Figure 1.**
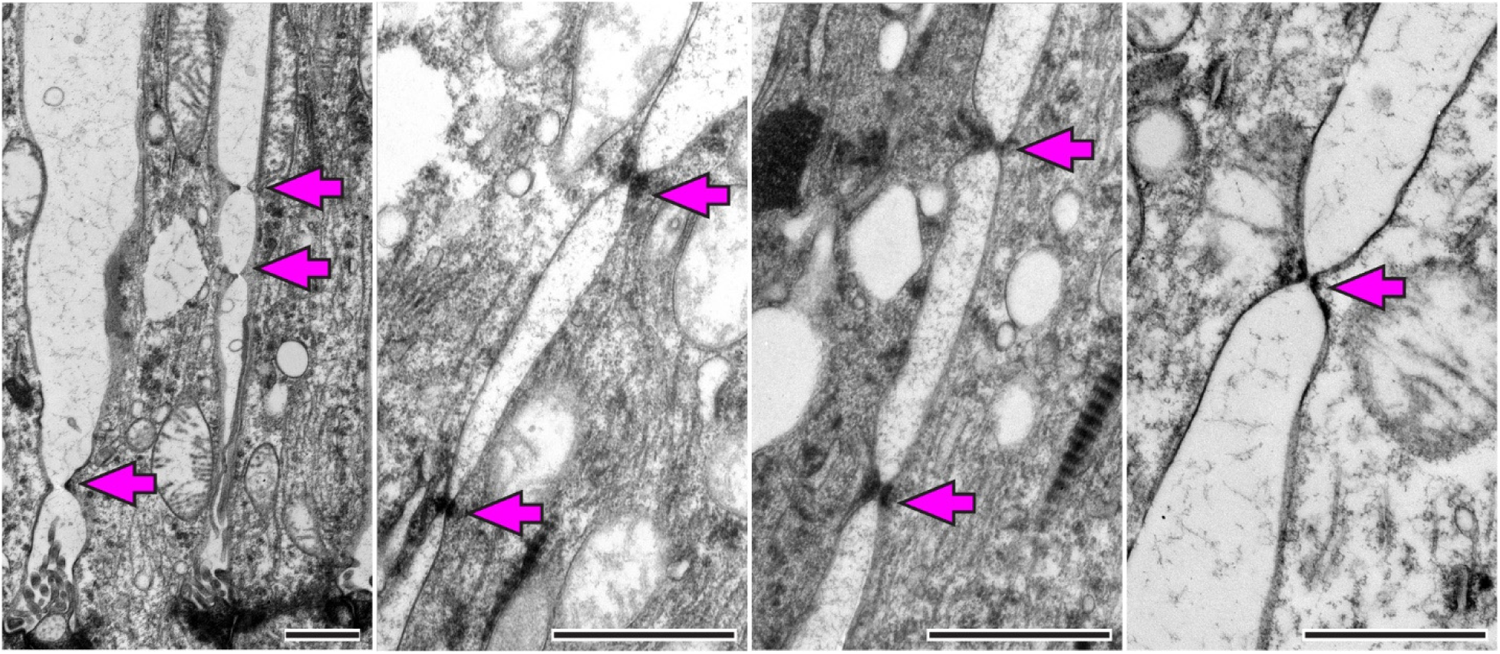
Human photoreceptors are connected by gap junctions. TEM images of photoreceptors from longitudinally sectioned human retinas. Magenta arrows indicate gap junctions connecting the IS of photoreceptor cells. Scale bars: 1 µm.

**Supplementary Figure 2.**
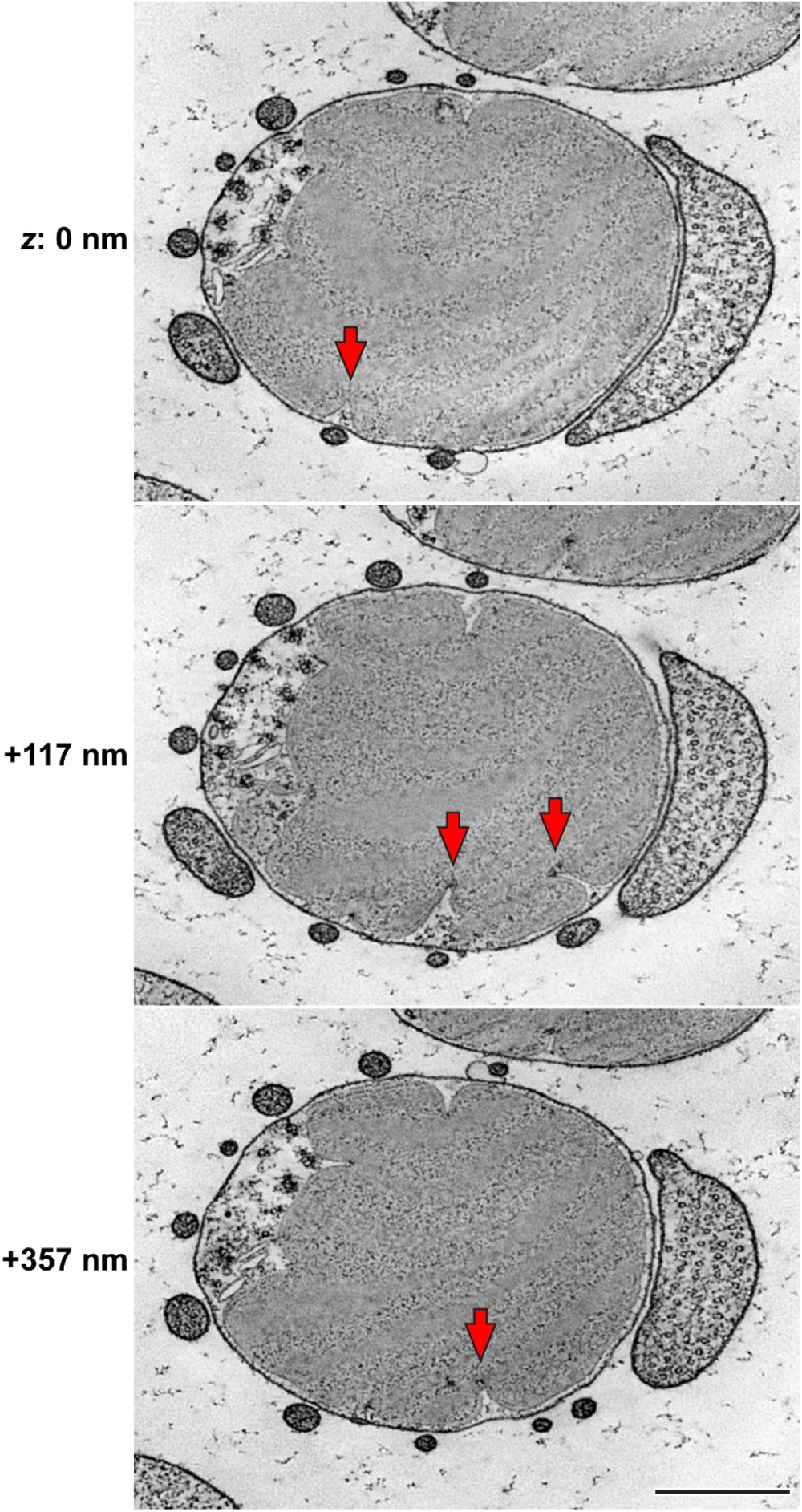
Disc incisures of human rods are not perfectly aligned. *z*-sections at the depths of 0 (top), +117 (middle) and +357 nm (bottom) from the tomogram in **Supplementary Video 1** illustrating an imperfect alignment of incisures across discs within the outer segment. Red arrows indicate incisures that are not aligned across the discs. Isotropic pixel size is 3 nm; scale bar: 0.5 µm.

**Supplementary Figure 3.**
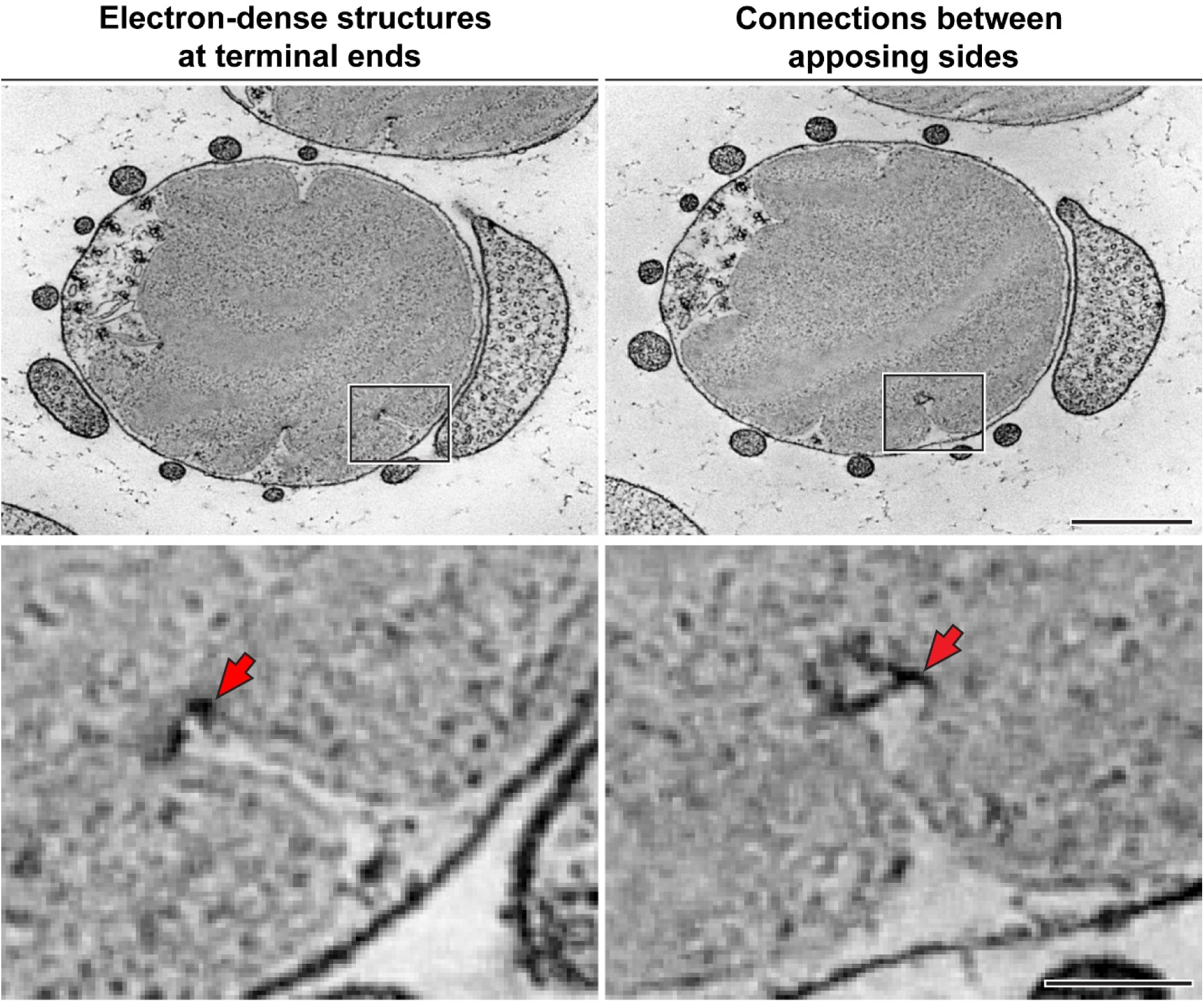
Ultrastructural features of human disc incisures. Maximum intensity projections of either 3 (left) or 4 (right) *z*-sections at different *z*-positions from the tomogram in **Supplementary Video 1** highlighting different ultrastructural features of human disc incisures: electron-dense structures at the terminal end (left) and connections between the apposing sides of an incisure (right). Red arrows indicate each corresponding feature of the incisure. Isotropic pixel size is 3 nm; scale bars: 0.5 µm (top); 0.1 µm (bottom).

**Supplementary Figure 4.**
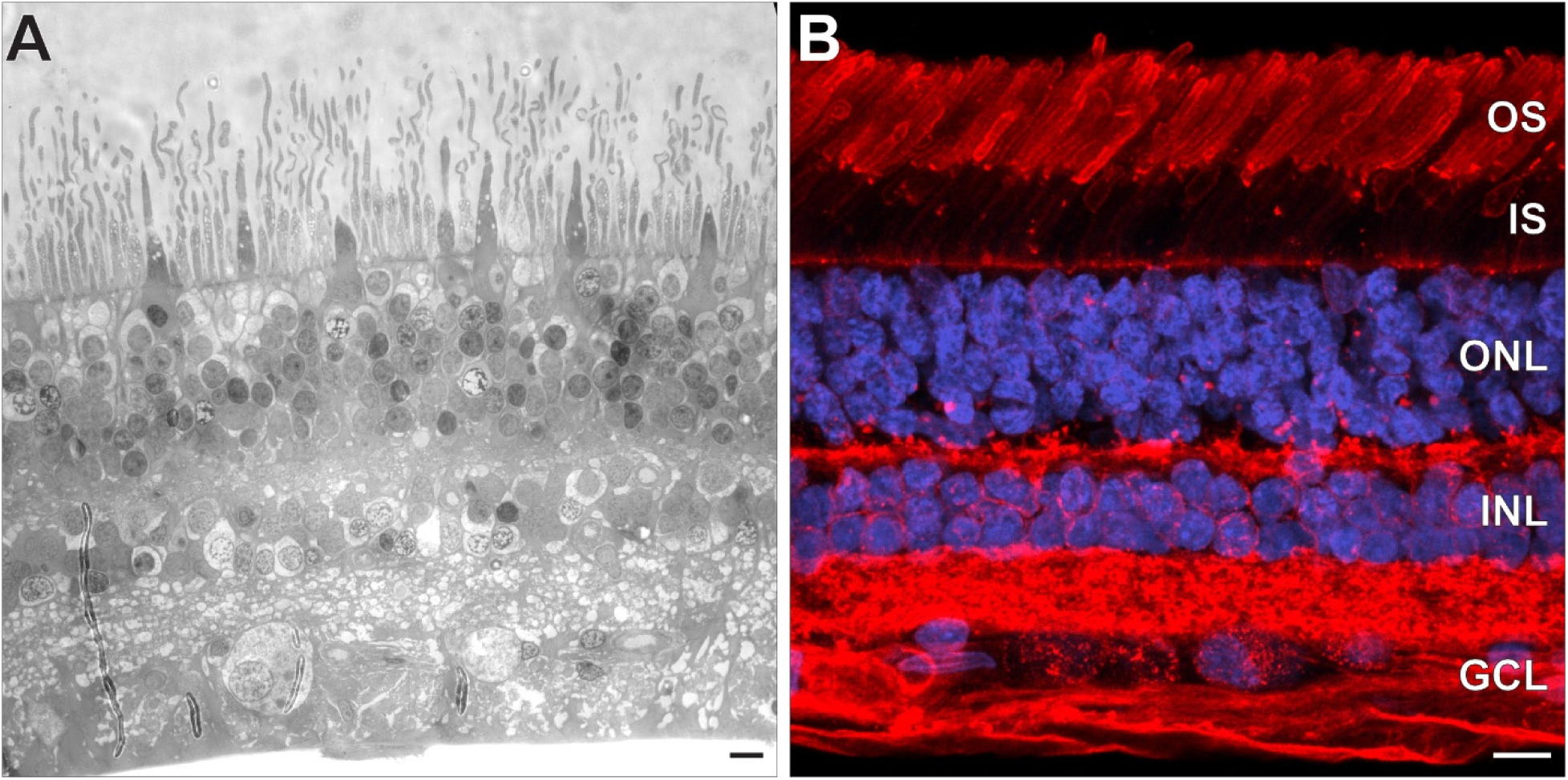
Tissue integrity of the human retina samples used in this study Bright-field. **(A)** and fluorescence **(B)** images of longitudinally sectioned human retinas illustrating the tissue preservation of the samples used for the ultrastructural and immunofluorescence analysis, respectively. WGA staining is shown in red, Höechst staining in blue. OS: outer segments; IS: inner segments; ONL: outer nuclear layer; INL: inner nuclear layer; GCL: ganglion cell layer. Scale bars: 10 µm.

